# PalmaClust: A graph-fusion framework leveraging the Palma ratio for robust ultra-rare cell type detection in scRNA-seq data

**DOI:** 10.64898/2026.03.16.712161

**Authors:** Xingzhi Niu, Jieqiong Wang, Shibiao Wan

## Abstract

**Motivation:** Single-cell RNA sequencing (scRNA-seq) is routinely used to build atlases of tissues, resolve developmental trajectories, and characterize disease microenvironments. Yet many biologically and clinically meaningful populations—including transient progenitors, therapy-resistant tumor subclones, and antigen-specific lymphocytes—occur at very low frequencies (*<*1%) and are easily missed by standard clustering pipelines. Existing approaches often require extensive manual curation, rely on known marker genes, or trade sensitivity for unacceptable false positive rates due to the insensitivity of metrics like the Gini index to heavy-tailed distributions. A scalable, statistically grounded method is needed to sensitively detect rare populations while providing calibrated confidence and interpretable molecular signatures.

**Results:** We present PalmaClust, a graph-fusion clustering framework that repurposes Palma ratio—a tail-sensitive inequality metric in sociology—to identify marker genes driven by extreme sparsity. PalmaClust constructs and fuses multiple K-Nearest Neighbor (KNN) graphs derived from complementary gene-selection statistics including the Palma ratio, Gini index, and Fano factor. It employs a local refinement strategy that re-prioritizes Palma-ranked genes within parent clusters. Benchmarking across diverse public scRNA-seq datasets confirms that PalmaClust consistently outperforms state-of-the-art baselines, improving rare-class F1 scores by at least 20% (absolute) while maintaining high global clustering stability. Further studies demonstrate that the Palma ratio-derived graph layer is essential for capturing ultra-rare signatures that other views miss.

**Availability:** https://github.com/wan-mlab/PalmaClust.

## 1. Introduction

Single-cell RNA sequencing (scRNA-seq) has fundamentally transformed our resolution of biological heterogeneity, yet the reliable detection of ultra-rare cell types, which often comprising less than 1% of a population, remains a formidable computational bottleneck (Kiselev *et al*., 2019). Detecting rare cell subpopulations is essential because rare populations often drive the most significant pathological and developmental processes. For instance, in acute myeloid leukemia (AML), primitive leukemic stem cell subpopulations often exhibit transcriptomic profiles that overlap significantly with background noise or unlabeled debris in standard annotations; failure to distinguish these rare subsets from technical artifacts obscures the origins of chemorefractory clones (Raffel *et al*., 2022). In solid tumor evolution, rare cells are the engines of progression: cancer stem cells (CSCs) (Trumpp and Haas, 2022), invasive mesenchymal subclones (Puram *et al*., 2017), and drug-tolerant persister cells (Sharma *et al*., 2010) often exist as minute populations that survive therapy to drive relapse and metastasis. Similarly, in the emerging field of liquid biopsy, circulating tumor cells (CTCs) represent ultra-rare events in blood samples where high contamination rates typically mask the signal of metastatic seeds (Andree *et al*., 2015). Furthermore, in cardiovascular research, endothelial progenitor cells (EPCs) have emerged as reliable biomarkers for vascular repair and aging, yet their scarcity in peripheral blood makes them difficult to quantify without highly sensitive, noise-robust methodologies (Williamson *et al*., 2012).

Standard clustering pipelines, which typically rely on variance-stabilizing transformations or the Fano factor, prioritize global structural variance. Consequently, they frequently fail to distinguish rare, biologically distinct signals from technical noise or ambient RNA contamination, leading to the merging of rare cells into major clusters or their dismissal as outliers (Young and Behjati, 2020). Current methods attempting to address this “needle-in-a-haystack” problem, such as GiniClust (Jiang *et al*., 2016) and the latest version GiniClust3 (Dong and Yuan, 2020) and RaceID (Grün *et al*., 2015) or its modern version RaceID3 (Herman *et al*., 2018), utilize the Gini index (Ceriani and Verme, 2011) or outlier probability models. However, the Gini index suffers from a well-documented economic limitation that translates directly to transcriptomics: it is most sensitive to the middle of the distribution and relatively insensitive to the tails (Gastwirth, 2017). In scRNA-seq, this means the signal of rare markers is often diluted by the moderate expression of housekeeping genes (Kiselev *et al*., 2019). Seurat (Satija *et al*., 2015) is a general-purpose scRNA-seq workflow that clusters cells via highly variable gene selection, principal component analysis (PCA), and community detection, but because it is optimized for global manifold structure, it often merges ultra-rare populations into nearby major clusters. ScCAD (Xu *et al*., 2024) frames rare-cell discovery as anomaly detection through cluster decomposition to isolate atypical substructures, but its performance can depend strongly on the initial representation/clustering and batch correction, potentially flagging transitional states or batch-driven artifacts as anomalies when rare signals are subtle.

To overcome these limitations, we present PalmaClust,a graph-fusion framework that repurposes the Palma ratio (Palma, 2011) for single-cell analysis. Originally defined in economics as the share of income held by the top 10% divided by the bottom 40%, the Palma ratio explicitly excludes the stable “middle”, acting as a high-pass filter for heavy-tailed distributions characteristic of rare markers (Mallick *et al*., 2022). Also, the Palma ratio offers a tunable mathematical framework (*p*_*top,bottom*_). Unlike the rigid Gini coefficient, the Palma calculation can be adapted to the expected sparsity of the target; for example, the numerator can be adjusted to focus on the top 1% or 5% to sensitize the metric for specific “ultra-rare” or “rare” priors, or “modified Palma ratios” can be employed to target specific distribution extremities. PalmaClust integrates this tail-sensitive logic within a multi-view graph learning architecture. We construct and fuse consensus K-Nearest Neighbor (KNN) graphs derived from complementary feature sets: Palma ratio, Gini index, and Fano factor, to capture both global structure and local rare-cell distinctiveness. By leveraging the Palma ratio’s unique sensitivity to extreme inequality, we demonstrated that PalmaClust significantly outperformed state-of-the-art approaches like GiniClust, RaceID, scCAD, and Seurat in recovering simulated and real-world rare populations, offering a robust solution for uncovering critical, low-frequency biological insights.

### 2. Materials and methods

### 2.1 The PalmaClust framework overview

Single-cell RNA-seq clustering methods often optimize global separation between major populations, which can obscure ultra-rare cell types whose marker genes are expressed in only a small fraction of cells. PalmaClust is designed to improve sensitivity to detecting these rare populations without sacrificing overall clustering quality. The overall workflow is shown in **Fig. 1**. Given a gene-by-cell count matrix, PalmaClust (a) scored genes with distribution-sensitive statistics, (b) constructed multiple complementary cell-cell KNN graphs from metric-specific feature sets, and fuses these graphs into a consensus “Mixed” graph, and (c) performed graph clustering followed by local refinement to resolve rare subtypes that may be merged at the global level.

**Fig. 1.**
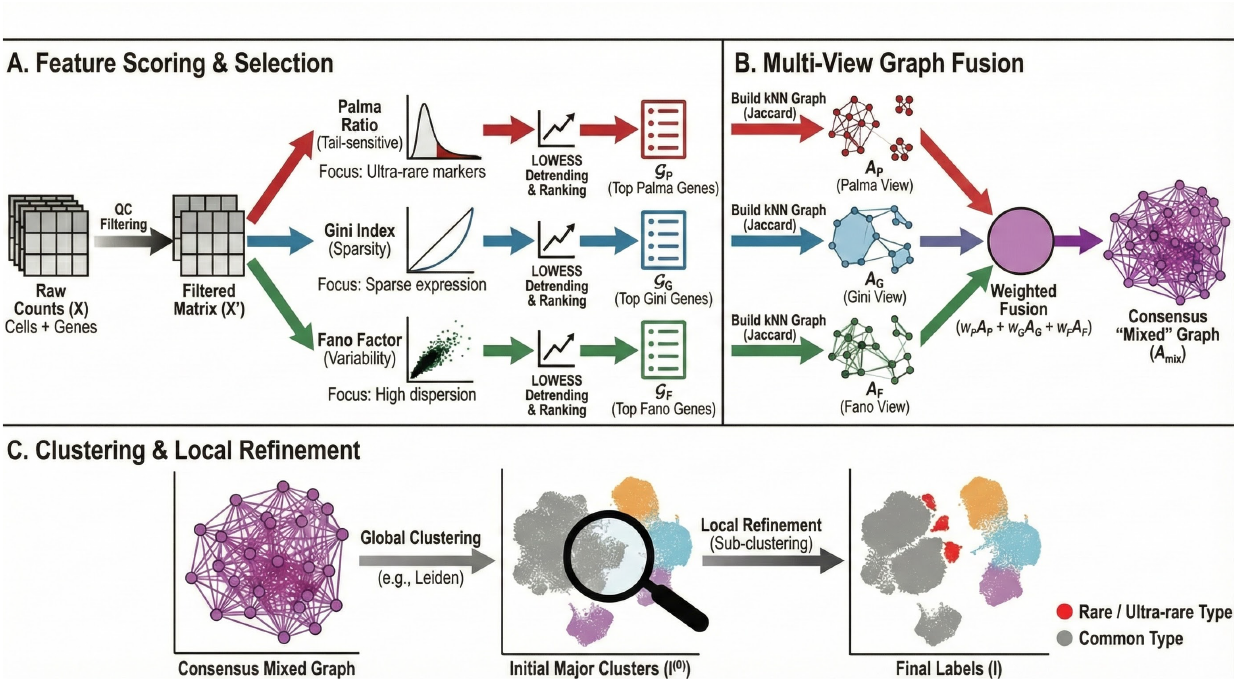
Overview of the PalmaClust pipeline for ultra-rare cell type detection. **(A) Feature scoring and selection**. Starting from a single cell count matrix **X**, quality control (QC) yields a filtered matrix **X**′. PalmaClust scores genes using three complementary statistics: the Palma ratio (tail-sensitive; enriches ultra-rare marker patterns), the Gini index (expression inequality/sparsity), and the Fano factor (dispersion/variability). Each score is LOWESS-detrended to reduce mean-dependent bias, then genes are ranked to produce three feature sets *G*_*P*_, *G*_*G*_, and *G*_*F*_ (top Palma/Gini/Fano genes).(**B**) **Multi-view graph construction and fusion**. Using each feature set, PalmaClust builds a metric-specific cell–cell kNN graph using Jaccard-based similarity (default), yielding three adjacency matrices **A**_*P*_ (Palma view), **A**_*G*_ (Gini view), and **A**_*F*_ (Fano view). These graphs are combined by weighted fusion, **A**_mix_ = *w*_*p*_**A**_*P*_ + *w*_*g*_ **A**_*G*_ + *w*_*f*_ **A**_*F*_, to form a consensus mixed graph that preserves global structure while strengthening rare-cell neighborhoods.(**C**) **Clustering and local refinement**. Community detection (Leiden) on *A*_mix_ produces initial major clusters 𝕀^(0)^, which are subsequently refined within clusters to resolve rare/ultra-rare subpopulations and output final labels 𝕀.

### 2.2 Preprocessing

Let **X** ∈ ℕ^*n*×*p*^ denote the input scRNA-seq count matrix with *n* cells and *p* genes, where *x*_*ig*_ is the observed UMI/read count of gene *g* in cell *i*. To reduce noise from extremely low-coverage cells and uninformative genes, we applied standard quality control filters prior to computing gene scores. Following the filtering strategy used in GiniClust-like pipelines, we excluded:

#### Low-coverage cells

remove cell *i* if the number of detected genes

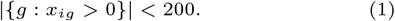

#### Rarely detected genes

remove gene *g* if it is expressed in fewer than 3 cells

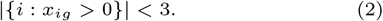

These thresholds (200 detected genes per cell; 3 expressing cells per gene) were treated as hyperparameters and can be tuned per dataset, but were fixed across methods in our benchmarks unless otherwise stated.

After filtering, we obtained a reduced matrix 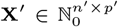 (with *n*′ ≤ *n, p*′ ≤ *p*). All downstream computations: gene scoring (Palma ratio/Gini index/Fano factor), feature selection, graph construction, graph mixing, and refinement, are performed on **X**′.

For each gene *g*, let **x**_*g*_ = (*x*_1*g*_, …, *x*_*n*_′ _*g*_) denote its expression across cells after filtering, and for each cell *i*, let **x**_*i*_ = (*x*_*i*1_, …, *x*_*ip*_′) denote its expression file. When computations require ordering expression values for a given gene, we wrote the sorted value as

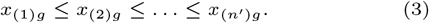

Unless explicitly stated, the metric definitions in subsequent sections were computed directly from the filtered raw counts **X**′, consistent with the empirically best-performing configuration in our experiments.

### 2.3 Gene scoring

Ultra-rare marker genes typically show heavy-tailed expression across cells: near-zero in most cells but high in a small subset. To explicitly emphasize this pattern, PalmaClust used a generalized Palma ratio-style ratio with tunable upper and lower fractions.

For each gene *g*, sort cells by expression *x*_*ig*_ in nondecreasing order:

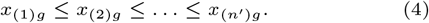

We defined two hyperparameters: (1) *p*_*t*_ ∈ (0, 1): the top fraction of cells, and (2) *p*_*b*_ ∈ (0, 1): the bottom fraction of cells, with the typical Palma ratio setting corresponding to (*p*_*t*_, *p*_*b*_) =(0.1, 0.4) in sociology, but both are adjustable.

Let 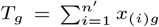 be the total expression mass. Define the top expression mass and the bottom expression mass as:

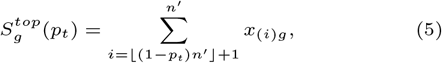

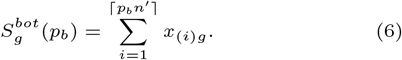

The generalized Palma score is then

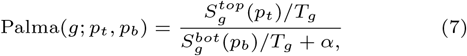

where *α* > 0 is a small constant to stabilize the denominator for sparse genes when the bottom mass is close to zero. In our experiment we constantly set *α* = 1e-6.

Intuitively, Palma(*g*; *p*_*t*_, *p*_*b*_) increases when a gene’s expression is concentrated in a small upper tail while remaining low in the lower portion of the population. This case is precisely the regime expected for ultra-rare markers. Decreasing *p*_*t*_ makes the score more sensitive to the most extreme tail (stronger rare emphasis), while increasing *p*_*b*_ increases denominator support and can improve stability in very sparse settings. In a real-world scenario, we may not know the rarity of the unknown population *a priori*. In our experiment, (*p*_*t*_, *p*_*b*_) = (0.1, 0.8) offered a stable bootstrap which works perfectly for cell types with rarity between 1% and 0.1%. We fixed that two parameters during all the following experiments.

To preserve global structure and provide complementary views of variability/inequality, PalmaClust also computed two standard gene-wise scores:

**Fano factor** (variance-to-mean ratio):

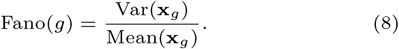

**Gini index** (expression inequality). With *µ*_*g*_ = Mean(**x**_*g*_) and sorted values *x*_(*i*)*g*_, one standard form is:

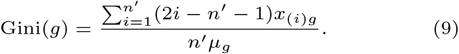

### 2.4 Feature selection

For each metric *m* ∈ {Palma(·; *p*_*t*_, *p*_*b*_), Gini, Fano}, we first computed a raw per-gene score 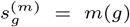. We observed that statistics can exhibit systematic dependence on a gene’s expression level, *e*.*g*., a gene with lower *log*_2_*max*(**x**_*g*_) tending to receive larger raw scores, PalmaClust applied a LOWESS (Seabold and Perktold, 2010) detrending step before ranking genes.

Concretely, let *M*_*g*_ = log_2_ max(**x**_*g*_) denote the logarithm of maximum expression of gene *g* from the filtered raw counts. For each metric *m*, we fitted a LOWESS curve

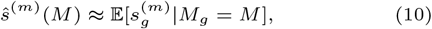

and computed the detrended score as residuals

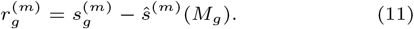

Genes were ranked by 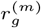 instead of raw 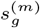 and the top *M* (*m*) genes were retained to define three feature sets: 𝒢_Palma_, 𝒢_Gini_, and *G*_Fano_, or correspondingly simplified as 𝒢_*P*_, 𝒢_*G*_, and 𝒢_*F*_.

Here the number of genes selected in the feature sets *M* (*m*) can be adjusted to optimize the performance, and in our experiment it was fixed to *M* (·) = 1000 as an acceptable robust bootstrap. These metric-specific gene sets were then used to construct complementary KNN graphs over cells, with Palma ratio emphasizing tail-driven rare-marker neighborhoods and Gini index and Fano factor contributing broader structural stability.

### 2.5 KNN graph construction

For each metric-specific gene set 𝒢_*m*_, *m* ∈ {Palma, Gini, fano}, PalmaClust constructed a cell-cell KNN graph *G*_*m*_.

Let **X**′ ∈ ℕ^*n*×*p*^ be the filtered raw count matrix. For a given metric *m*, we extracted the selected feature submatrix 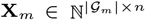 containing only genes in *G*_*m*_. We then determined a single binarization cutoff

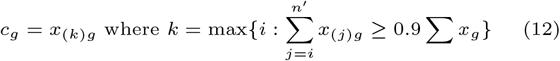

for each gene, *i*.*e*. the lowest value included in the top 90% of the total summation. Using the cutoff, we activated each gene in each cell as

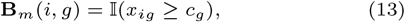

yielding a sparse binary matrix 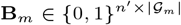.

Given binary feature vectors **b**_*i*_ and **b**_*j*_ for cells *i* and *j* which are rows of **B**_*m*_, we computed Jaccard similarity:

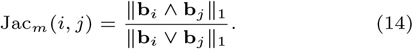

For each cell *i*, we selected its *K* = 30 nearest neighbors under maximum Jaccard similarity, forming a directed KNN relation that is then symmetrized by edge union to obtain an undirected graph *G*_*m*_ with adjacency **A**_*m*_. Edge weights were set to:

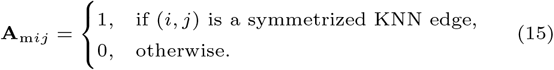

Applying this procedure to 𝒢_*P*_, 𝒢_*G*_, 𝒢_*F*_ yielded three complementary graphs: *G*_Palma_, *G*_Gini_, and *G*_Fano_, or simplified as *G*_*P*_, *G*_*G*_, and *G*_*F*_.

### 2.6 Graph mixing and initial clustering

PalmaClust integrated the three complementary KNN graphs *G*_*P*_, *G*_*G*_, and *G*_*F*_ into a single consensus graph that simultaneously preserved global population structure and enhanced rare-cell separability.

Let 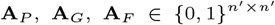 denote the symmetrized adjacency matrices for the three graphs. PalmaClust forms the mixed adjacency matrix:

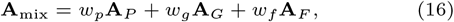

where *w*_*p*_, *w*_*g*_, *w*_*f*_ *≥* 0 and *w*_*p*_ + *w*_*g*_ + *w*_*f*_ = 1.

Intuitively, **A**_*P*_ tends to reinforce local neighborhoods driven by tail-dominant, *i*.*e*., rare-marker genes, while **A**_*G*_ and **A**_*F*_ contribute stabilizing signal for abundant populations and overall manifold structure. The convex combination above enabled PalmaClust to balance these complementary objectives via (*w*_*p*_, *w*_*g*_, *w*_*f*_). As for general applications, (*w*_*p*_, *w*_*g*_, *w*_*f*_) = (0.5, 0.1, 0.4) offers a robust bootstrap and was used for all following experiments unless explicitly stated otherwise.

We obtained the initial clustering by applying a graph community detection algorithm (*e*.*g*. in our experiments, Leiden (Traag *et al*., 2019)) to the mixed graph defined by **A**_mix_. This produces an initial partition of the cells into communities {**C**_1_, …, **C**_*K*_ }, which then served as the starting point for the local refinement step.

### 2.7 Local refinement for discovering rare and ultra-rare subpopulations

After obtaining an initial partition from the mixed global graph, PalmaClust performed a within-cluster refinement step to identify rare substructures that may be diluted at the whole-dataset scale. Refinement was applied independently inside each parent (major) cluster **C**.

For each parent cluster **C** and a chosen refinement gene pool *G* (in our experiments, 𝒢 = 𝒢_Palma_, *i*.*e*., we reused the genes selected by Palma ratio), we extracted the local matrix **X**_𝒢,**C**_ from the filtered raw counts. We then constructed a local similarity graph among cells in **C** using: (1) low dimensional embedding: applied truncated singular value decomposition (SVD) to obtain a dense cell embedding by taking first 20 values, and (2) Cosine KNN graph: a cosine-similarity KNN graph with *K* = 30 **A**_**C**_ was built from the truncated SVD features. PalmaClust hybridized this local graph with the corresponding subgraph of the global mixed graph restricted to **C**:

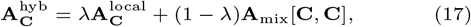

where *λ* ∈ [0, 1] is a hyperparameter. This improved robustness when rare markers are weak but the global manifold structure is reliable. In our experiment we set *λ* = 0.7.

## 3. Results

### 3.1 Palma ratio prioritized ultra-rare cell type markers

To validate that PalmaClust’s feature scoring is intrinsically sensitive to rare signals rather than relying solely on downstream graph mixing, we tested whether different gene-ranking metrics preferentially elevate known rare-type markers under controlled rarity. We diluted the rare cell type ionocytes (29 out of 14,163, *≈* 0.205%) in GSE102580 (Plasschaert *et al*., 2018) by generating background cells with negative binomial dispersion. Using GSE102580-derived augmented datasets, we progressively diluted the target rare population by expanding the non-target background while keeping the marker set fixed (top 10 Wilcoxon markers computed once on the original dataset). For each augmented dataset, we computed Palma ratio-, Fano factor-, and Gini index-based gene scores with the same LOWESS detrending procedure, ranked genes genome-wide, and evaluated where the marker genes appeared in the ranked list.

Across all dilution proportions, the Palma ratio consistently ranked rare-type markers closer to the top. As shown in **Fig. 2**, Palma ratio achieved the highest average marker rank percentile, remaining around the low-to-mid 90th percentiles throughout the dilution range. In comparison, Fano factor ranked the same markers slightly lower (*≈*86–92nd percentiles) and decayed with the rare type being further diluted, while Gini index exhibited substantially poorer prioritization (*≈*78–85th percentiles) and greater variability across dilution settings.

**Fig. 2.**
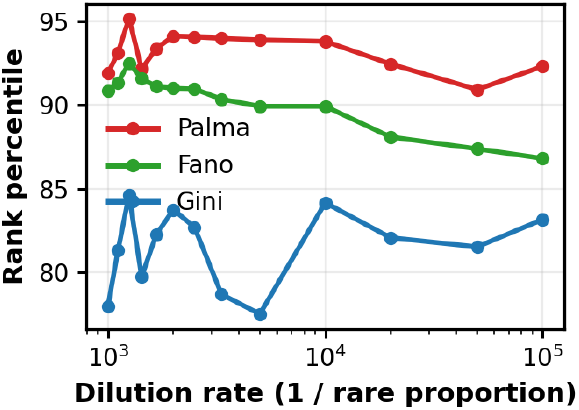
Palma ratio robustly prioritized rare-cell marker genes under progressive dilution of the rare population. Average marker rank percentile (higher is better) for a fixed marker set (*N* = 10), computed once from the original dataset using a Wilcoxon rank-sum test (top 10 markers) and held constant across all augmented (diluted) datasets. Genes were ranked by the corresponding (LOWESS-detrended) feature scores from Palma ratio, Fano factor, or Gini index (shown as “Palma”, “Fano”, and “Gini” in the figure correspondingly). Across dilution rates (x-axis: 1*/*rare proportion), Palma ratio consistently assigned the highest average percentile to the marker set.

This indicated that Palma more reliably prioritizes ultra-rare marker genes upward in the global ranking as the target population becomes increasingly scarce. This results supported PalmaClust’s central design choice: Palma-based scoring provides robust rare-marker prioritization even when the rare population frequency is pushed into an ultra-rare regime, forming a stronger foundation for subsequent graph construction and rare-cell separability.

### 3.2 Graph fusion enhanced separability of rare cells

To assess whether PalmaClust’s improvements arise at the graph-geometry level before clustering, we compared how different feature-selection metrics shape the cell-cell neighborhood structure. For GSE102580, we constructed metric-specific kNN graphs using gene sets selected by Fano factor (*G*_Fano_), Gini index (*G*_Gini_), and Palma ratio (*G*_Palma_), and then formed the fused consensus graph (*G*_mix_) by weighted combination of these adjacency matrices. The method to generate the graphs was identical to the method of PalmaClust pipeline. To visualize how each graph organizes cells, we computed t-SNE embeddings directly from the graph-derived distance structure (precomputed distances from the kNN adjacency matrix), highlighting the target ultra-rare ionocytes. As shown in **Fig. 3A–B**, both *G*_Fano_ and *G*_Gini_ yielded embeddings where ionocytes were split into multiple locations rather than forming a stable, isolated neighborhood. This fragmentation suggested that dispersion- or sparsity-based features alone may not reliably reinforce rare-to-rare edges when the rare population is extremely small, causing rare cells to attach to distinct major clusters in the kNN graph. In contrast, *G*_Palma_ produced a substantially more coherent arrangement (**Fig. 3C**), consistent with Palma’s tail-sensitive ranking elevating genes whose expression mass is concentrated in a small subset of cells, thereby strengthening rare-cell connectivity. Nevertheless, *G*_Palma_ itself could not separate major types well because it omits connections between major type cells.

**Fig. 3.**
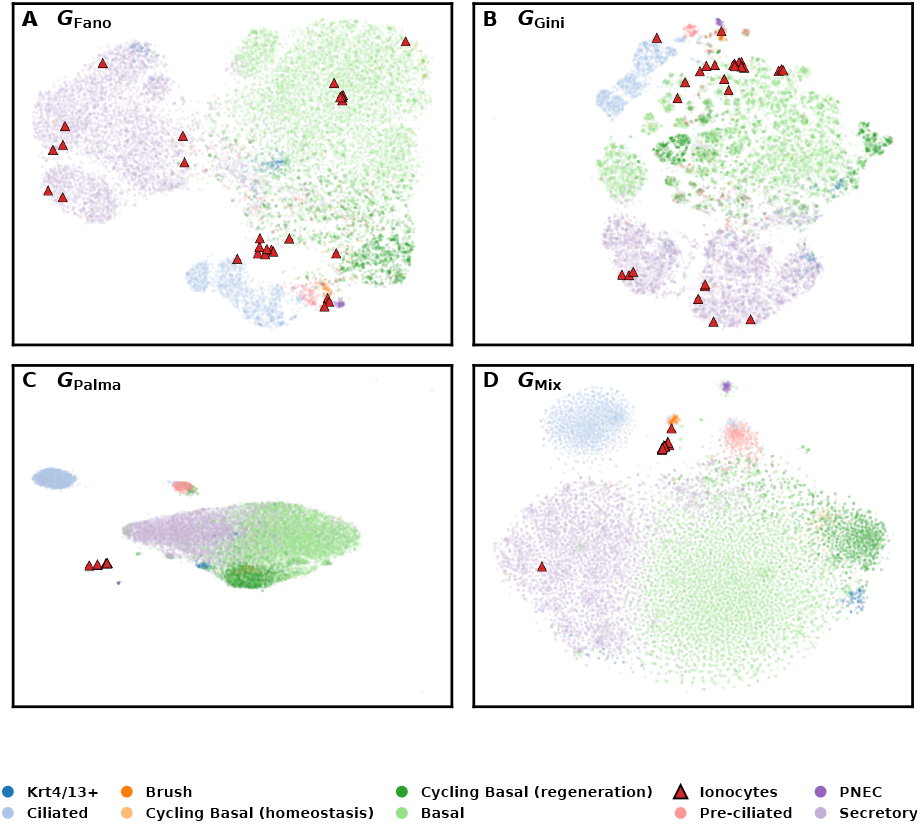
Graph-dependent rare-cell separability visualized by t-SNE from kNN adjacency structure for GSE102580. t-SNE embeddings are computed using precomputed cell-cell distances derived directly from the kNN adjacency matrix of each graph view from PalmaClust pipeline of GSE102580. Cells are colored by reference cell type labels, and the target ultra-rare population (Ionocytes) is highlighted as red triangles. **(A)** *G*_Fano_: ionocytes are dispersed across multiple regions, indicating that the variability-driven graph connects rare cells to different major neighborhoods rather than forming a coherent rare-cell community.**(B)** *G*_Gini_: ionocytes remain fragmented and partially mixed with major populations, suggesting inequality alone is insufficient to stabilize ultra-rare neighborhoods. **(C)** *G*_Palma_: ionocytes become more locally coherent, consistent with Palma’s tail-sensitive gene selection producing a graph that strengthens rare-cell-to-rare-cell connectivity. However due to the involvement of very tail-sensitive signals, which could be noise to the global manifold structure, the major types are not well separated. **(D)** *G*_mix_: the fused graph yields the most concentrated ionocyte neighborhood while maintaining the global separation of major cell groups, illustrating that multi-view fusion preserves overall structure while enhancing rare-cell separability.

Finally, the fused graph *G*_mix_ provides the clearest qualitative separation (**Fig. 3D**): ionocytes formed a tight and consistent neighborhood while the major epithelial populations remain well organized. This supported the central hypothesis of PalmaClust: Palma ratio-derived structure is critical for rare-cell cohesion, but combining Palma ratio with complementary views (Fano factor/Gini index) helps preserve global manifold structure. This qualitative improvement in rare-cell separability anticipated the quantitative gains in rare-type detection reported in subsequent benchmarking analyses.

### 3.3 PalmaClust detected rare cell types while preserving major populations clustering performance

We first benchmarked PalmaClust with GSE102580, a single-cell atlas of the airway epithelium that reported the CFTR-rich pulmonary ionocyte as a newly characterized epithelial cell type. Ionocytes are biologically important because they are enriched for CFTR and ion-transport programs (e.g., FOXI1 and V-ATPase components), linking this population to airway physiology and cystic fibrosis–relevant mechanisms. Importantly for method evaluation, ionocytes are also extremely rare, making them an archetypal “needle-in-a-haystack” clustering target.

After QC filtering, our GSE102580 analysis set contained 14,163 cells spanning major epithelial populations (Basal: 6,009; Secretory: 4,792; Ciliated: 1,333; Cycling Basal regeneration: 1,242; *etc*.) and several low-frequency types, including Ionocytes: 29 cells (0.2%). This prevalence (*≈* 2 × 10^−3^) placed ionocytes firmly in the ultra-rare regime, where global-structure-driven clustering objectives often merge them into abundant neighbors. For all benchmarkings, we took the same filtering step for baseline methods as PalmaClust. We tuned the clustering parameters to the best performance of GiniClust, Seurat, and RaceID3. Due to the blackbox design of GiniClust3 and scCAD implementation, we take the recommended default setup of them.

As shown in **Fig. 4A**, PalmaClust recovered ionocytes with F1 = 0.87, exceeding RaceID3 (0.65) and scCAD (0.59) by a wide margin, and GiniClust (F1 *<*0.01) and GiniClust3 (F1 *<*0.01) nearly failed to identify ionocytes, consistent with the ionocyte signal being absorbed into major epithelial clusters. The reason GiniClust and GiniClust3 failed to identify the ultra-rare cell type might be because Gini index was significantly influenced by the “middle”, which was consistent with **Fig. 2**. In **Fig. 4B**, PalmaClust achieved strong overall agreement with the reference labels (ARI = 0.74, NMI = 0.71), outperforming all baselines like RaceID3 and Seurat on ARI (0.16 and 0.28, respectively). Crucially, PalmaClust’s global performance did not come at the expense of rare sensitivity.

**Fig. 4.**
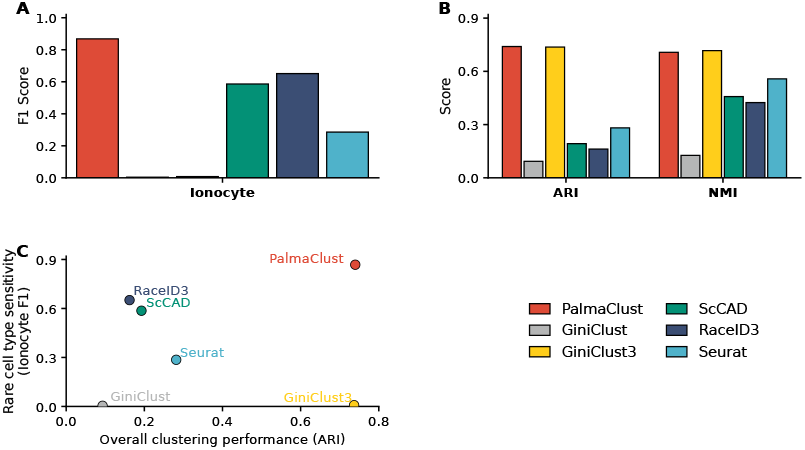
PalmaClust accurately identified the rare cell type ionocyte for the GSE102580 without sacrificing global clustering quality. **(A)** Rare cell type detection performance measured by F1 score for the ionocyte population (29 cells; 0.2%). PalmaClust yielded the highest rare-cell accuracy (F1 = 0.87), outperforming RaceID3 (0.65), scCAD (0.59) and Seurat (0.29), while GiniClust3 (*<*0.01) and GiniClust showed near-zero ionocyte recovery (*<*0.01). **(B)** Overall clustering performance measured by ARI and NMI against the reference labels. PalmaClust achieved strong global agreement (ARI = 0.74, NMI = 0.71), comparable to the best overall baseline (GiniClust3: ARI = 0.74, NMI = 0.72), and outperforming all other baselines (best of others was Seurat: ARI = 0.28, NMI = 0.56). **(C)** Trade-off between global clustering performance (ARI; x-axis) and rare-cell sensitivity (ionocyte F1; y-axis).

The joint view of global accuracy versus rare sensitivity (**Fig. 4C**; ARI vs ionocyte F1) highlights the central advantage of PalmaClust on this dataset: it was the only method that simultaneously attains high overall clustering quality and high ultra-rare recovery for a population comprising only 29 out of 14,163 cells.

We next evaluated PalmaClust with GSE94820 (Villani *et al*., 2017), a human blood myeloid scRNA-seq dataset associated with a single-cell RNA-seq study of new types of human blood dendritic cells, monocytes and progenitors.

This dataset is a strong benchmark for rare detection because it contains multiple closely related dendritic cell (DC) and monocyte subtypes with subtle transcriptional boundaries. In such settings, rare subtypes are easily merged into adjacent abundant populations unless the clustering representation explicitly amplifies tail/rarity signals.

After QC filtering, our GSE94820 analysis set contained 1,123 cells distributed across annotated DC and monocyte subtypes. While several populations were moderately sized (*e*.*g*., DC4: 182 cells, 16.2%; DC6: 173, 15.4%), the dataset included one low-frequency cell type (Mono4: 18 cells =1.6%) that were appropriate for rare-sensitivity evaluation.

On rare-subtype Mono4 recovery (**Fig. 5A**), PalmaClust achieved the best detection performance (Mono4 F1 = 0.78). In contrast, competing approaches showed markedly reduced sensitivity in this regime (scCAD: 0.36, RaceID3: 0.15, and Seurat: 0.15). Again GiniClust3 and GiniClust failed to identify the rare cell type because Gini index lost the sensitivity to rare cell types in the ~ 1% to 0.1% regime.

**Fig. 5.**
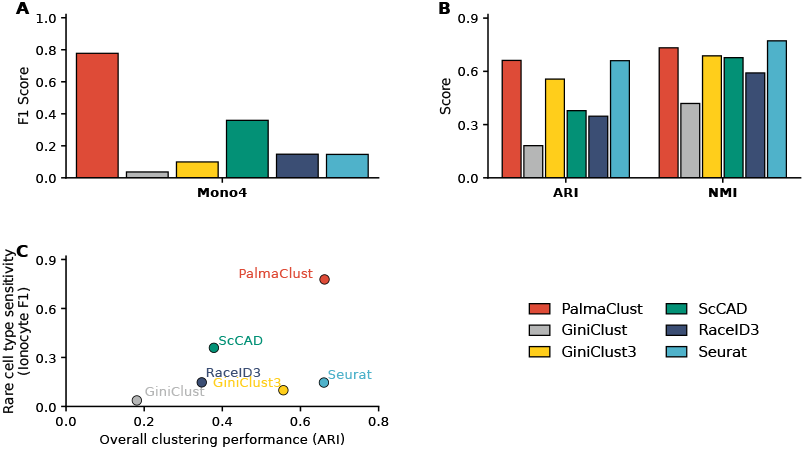
PalmaClust achieved strong rare cell type detection sensitivity for the GSE94820 while maintaining competitive overall clustering accuracy. **(A)** Rare cell type detection performance measured by F1 score for the low-frequency target populations in GSE94820: Mono4 (18 cells; 1.6%). PalmaClust achieved the highest F1 for Mono4 (0.78). It outperformed scCAD (0.36), RaceID3 (0.15), Seurat (0.15), GiniClust3 (0.10) and GiniClust (0.04). **(B)** Overall clustering performance on GSE94820 measured by ARI and NMI against the reference labels. PalmaClust attained high global agreement (ARI = 0.66, NMI = 0.73), comparable to the strongest overall-performing baseline (Seurat, ARI = 0.66, NMI = 0.77). **(C)** Trade-off between global clustering performance (ARI, x-axis) and rare sensitivity (Mono4 F1, y-axis).

On overall clustering quality (**Fig. 5B**), PalmaClust achieved ARI = 0.66 and NMI = 0.73, remaining competitive with the strongest global baseline (Seurat: ARI = 0.66, NMI = 0.77) and outperforming GiniClust3 (ARI = 0.56, NMI = 0.69), RaceID3 (ARI = 0.35, NMI = 0.59) and scCAD (ARI = 0.38, NMI = 0.68).

Finally, the trade-off plot (**Fig. 5C**; ARI vs Mono4 F1) illustrates that PalmaClust uniquely occupied the high-high region—simultaneously preserving the global DC/monocyte structure and isolating a 1.6% population (Mono4) as a coherent cluster—supporting the generalization of PalmaClust beyond airway epithelium to a challenging immune subtype landscape.

### 3.4 Palma ratio-based component was essential to rare cell type detection

To quantify how each graph view contributed to PalmaClust, we performed a systematic graph-fusion weight scan over the simplex (*w*_*f*_, *w*_*g*_, *w*_*p*_) with *w*_*f*_ + *w*_*g*_ + *w*_*p*_ = 1, where *w*_*f*_, *w*_*g*_, and *w*_*p*_ weighted the Fano factor, Gini index, and Palma ratio-based kNN graphs, respectively. For each weight triplet with step size 0.1, we ran the full PalmaClust pipeline and evaluated (i) global clustering agreement using ARI and (ii) ultra-rare detection accuracy using Ionocytes F1 on GSE102580, where ionocytes comprise only 29 cells (0.20%) of the atlas. The resulting heatmaps (**Fig. 6A–B**) showed that removing Palma (*w*_*p*_ = 0, the leftmost column) had a strong negative impact on both objectives: ARI drops substantially (**Fig. 6A**), and ionocyte detection collapsed to near-zero F1 (**Fig. 6B**). In contrast, mixtures that retained a non-zero Palma component yielded markedly stronger rare-cell sensitivity, with ionocyte F1 generally increasing as *w*_*p*_ increased, consistent with Palma-derived features strengthening rare-to-rare neighborhood connectivity.

**Fig. 6.**
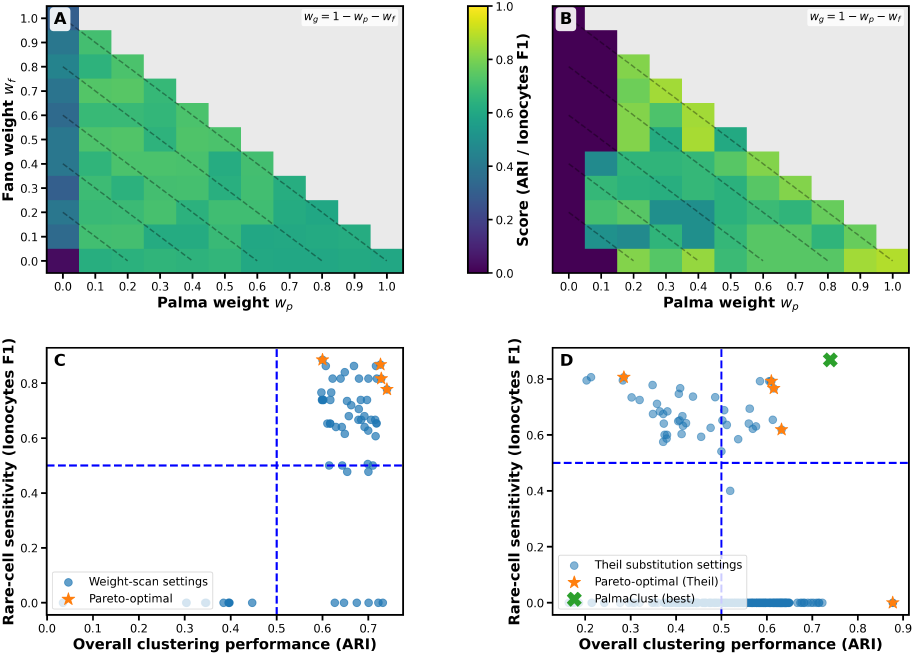
Ablation and sensitivity analysis of the graph-fusion mechanism highlighted the necessity of the Palma component for balancing global clustering accuracy and ultra-rare cell type detection (GSE102580; ionocytes: 29/14163 cells, 0.20%). **(A)**Heatmap of overall clustering performance (ARI) under a fusion-weight scan, visualized on a (*w*_*p*_, *w*_*f*_) grid with *w*_*g*_ = 1 − *w*_*p*_ − *w*_*f*_ (only feasible combinations with *w*_*p*_ + *w*_*f*_ ≤ 1 are shown). Removing the Palma component (*w*_*p*_ = 0, left edge) substantially degraded ARI, whereas balanced mixtures yielded the strongest global performance. Diagonal dashed lines denote iso-*w*_*g*_ (constant Gini weight). **(B)** Heatmap of rare-cell sensitivity (ionocyte F1) on the same weight grid. Without Palma (*w*_*p*_ = 0), ionocyte detection collapsed (near-zero F1), while increasing *w*_*p*_ generally improved ionocyte recovery, indicating that Palma-derived features are critical for forming rare-cell neighborhoods. **(C)** Pareto plot of the PalmaClust fusion-weight scan, showing the trade-off between global accuracy (ARI) and rare-cell sensitivity (ionocyte F1). Blue dashed lines indicate reference thresholds (0.5 for ARI and 0.5 for F1); orange stars mark Pareto-optimal settings. A representative balanced configuration (*w*_*f*_, *w*_*g*_, *w*_*p*_) = (0.5, 0.1, 0.4) achieved strong performance on both axes. **(D)** Pareto plot for a substitution ablation in which Palma scoring was replaced by the Theil index (with the same LOWESS detrending and fusion-weight tuning), compared against the PalmaClust best reference point (green cross; ARI=0.7398, ionocyte F1=0.8679). While some Theil settings achieve competitive ARI or F1 individually, the Pareto front failed to match PalmaClust’s combined performance.

This trade-off was summarized by the Pareto plot over the same scan (**Fig. 6C**), which directly visualized the achievable combinations of ARI and ionocyte F1. The blue dashed lines at 0.5*/*0.5 indicated a pragmatic acceptable region for both global clustering and rare sensitivity. Pareto-optimal solutions concentrated in the upper-right, confirming that PalmaClust can simultaneously achieve competitive ARI and strong ionocyte detection, but only when Palma contributed meaningfully to the fused graph.

We further tested whether Palma’s role could be replaced by other feature-scoring criteria under the same detrending and tuning protocol. First, we substituted Palma scoring with the Theil index, an inequality measure defined per gene *g* across *N* cells with expression *x*_*g*1_, …, *x*_*gN*_ and mean *µ*_*g*_ as

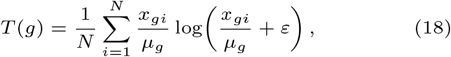

where *ε* = 10^−6^ avoids log(0). We ranked genes by *T* (*g*), applied the same LOWESS detrending, constructed the corresponding kNN graph, and repeated graph-fusion tuning. Although some Theil settings can achieve competitive ARI or competitive ionocyte F1 individually, they fail to maintain both simultaneously: the Theil substitution Pareto front remains far from the PalmaClust reference point (ARI= 0.7398, ionocyte F1= 0.8679) (**Fig. 6D**). We also evaluated two additional substitutions: (i) a 90% coverage score that measures concentration by the smallest number of cells *c*_0.9_(*g*) needed to account for 90% of a gene’s total expression mass (genes ranked higher when *c*_0.9_(*g*) is smaller), and (ii) an IDF score based on detection frequency 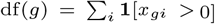 defined as 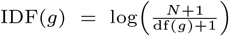. Neither replacement was successful: 90% coverage achieved a maximum ARI of 0.447 and no setting detected ionocytes (ionocyte F1*≈* 0), while IDF achieved a maximum ARI of 0.445 and likewise never detected ionocytes. Together, these ablations indicated that PalmaClust’s performance was not explained by generic sparsity or inequality scoring; rather, the Palma component was specifically required to achieve the key objective of high rare-cell sensitivity without sacrificing global clustering quality.

### 3.5 PalmaClust was scalable and computationally efficient

We next evaluated the computational scalability of PalmaClust and compared its runtime characteristics with widely used clustering/rare-cell detection baselines. All the time and memory benchmarking below were the average of three runs. We first compared end-to-end step breakdowns on the GSE102580 dataset (14,163 cells) across methods (**Fig. 7A**).PalmaClust completed in ~15 s total, with refinement being the largest component at this small scale, whereas Seurat required ~159 s, GiniClust required ~303 s, and GiniClust3 required ~197 s under the same single-threaded setting. ScCAD incurred substantial time in its combined compute stage (reported as an aggregated gene-selection/clustering block). RaceID3 was dominated by an extremely long clustering stage (~ 6.0×10^4^ s), resulting in a total runtime of ~17.3 h on the same dataset, highlighting its limited scalability for large-scale analyses. It is worth noting that PalmaClust reached the shortest execution time in every stage of rare cell type detection pipeline, and a low level C++ based I/O interface of PalmaClust allowed a significant reduction of reading and writing time.

**Fig. 7.**
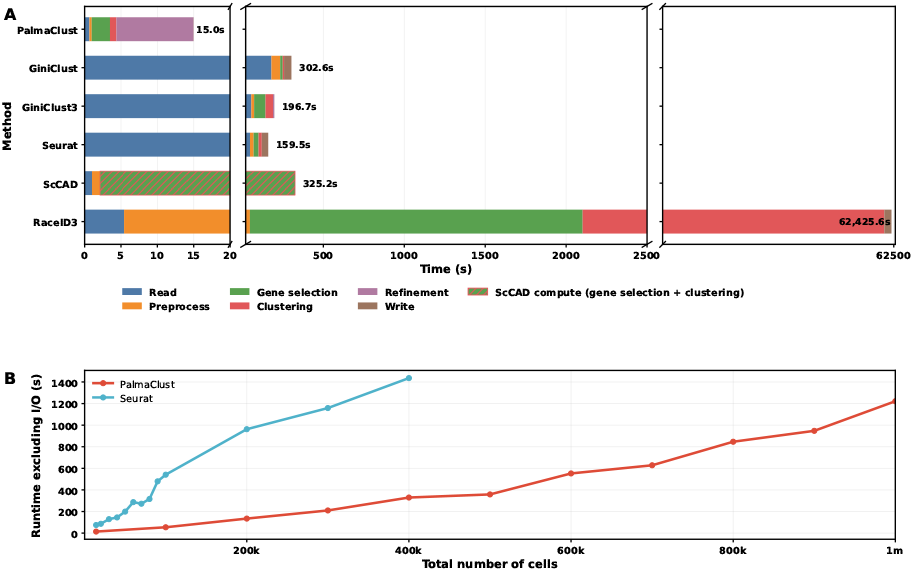
Scalability and runtime breakdown of PalmaClust and baselines. **(A)** Stepwise wall-time breakdown on GSE102580 (14,163 cells) using a stacked horizontal bar chart with two axis breaks to visualize both the sub-20 s regime and the long tail of RaceID3. Bars are decomposed into pipeline stages (read, preprocess, gene selection, clustering, refinement, write); totals are annotated at the bar ends. For scCAD, the reported compute time is available only as an aggregated stage and is shown as a hatched block to indicate combined gene-selection/clustering cost. All timings were obtained with single-threaded execution; PalmaClust additionally supported CUDA acceleration for kNN graph construction, which yielded > 10× speedup for that stage in profiling. **(B)** Runtime scaling (excluding I/O) as a function of total number of cells. PalmaClust scaled to > 2×10^6^ cells, while Seurat showed the most moderate growth among the tested baselines within the feasible regime but became much slower than PalmaClust beyond ~ 4 × 10^5^ cells in our setting.

We then performed a scaling test by increasing the number of cells up to 2 × 10^6^ by augmenting the GSE102580 dataset, measuring wall time excluding I/O. In this setting,PalmaClust completed successfully beyond 1M cells (**Fig. 7B**), demonstrating practical scalability to modern atlas-sized scRNA-seq datasets. In contrast, among the baseline methods we tested, Seurat exhibited the only relatively moderate growth curve within the feasible regime, and we therefore focused the direct growth comparison on Seurat; notably, Seurat became impractical in our environment beyond ~ 4 × 10^5^ cells due to rapidly increasing runtime and resource requirements (**Fig.7B**).

A key design choice enabling large-scale execution was that PalmaClust maintained sparsity throughout graph construction. After the activation/binarization step, the expression matrix remained highly sparse with an approximately constant density of ~ 0.4% nonzero entries (nnz) across scales. Furthermore, the cell–cell graph was explicitly pruned to a fixed top-*K* neighborhood, yielding an adjacency structure with *O*(*nK*) edges rather than *O*(*n*^2^). This sparsity and top-*K* pruning effectively control steps that would otherwise be quadratic in the number of cells when implemented with dense pairwise computations.

Stepwise profiling across increasing cell numbers further clarified where computation concentrates as scale grows. Empirically, the near-linear components include basic metric computation and binarization (approximately proportional to nnz), while detrending and gene selection remain comparatively small and weakly dependent on *n*. The dominant costs at large *n* are (i) KNN graph construction and fusion and (ii) community detection (Leiden), which increase superlinearly with cell number. Importantly, PalmaClust supported CUDA (Nickolls *et al*., 2008) acceleration for kNN graph construction;in our profiling this reduced the KNN stage runtime by more than an order of magnitude (> 10×) relative to a CPU-only implementation, substantially improving end-to-end scalability.All results reported here were executed in a single-threaded configuration; while we have not yet ported Leiden or the refinement passes to CUDA, these stages are also graph-based and therefore represent promising targets for further acceleration.

Finally, PalmaClust’s sparse-matrix implementation reduced memory usage compared with dense representations: on large samples (> 10^5^ cells) we observed > 2× lower memory footprint, consistent with the efficiency achieved by other sparse-aware toolchains such as Seurat. Together, these results indicated that PalmaClust combined rare-cell sensitivity with practical scalability, reaching multi-million-cell regimes while maintaining controlled compute and memory growth (Fig. 7A–B).

At a high level, PalmaClust’s pipeline can be understood as (i) sparse per-gene metric computation and activation (*≈ O*(nnz)), (ii) top-*K* KNN construction and weighted graph fusion (practically *O*(*nK*) edges, with KNN search cost dominating), (iii) community detection on a sparse graph (approximately linear in the number of edges in practice), and (iv) local refinement passes that operate on subgraphs and remain sparse. In contrast, methods whose bottlenecks involve expensive iterative clustering or implicit dense pairwise operations exhibited substantially worse empirical scaling, as reflected by RaceID3 in our benchmarks (**Fig. 7A**).

### 4. Discussions

### 4.1 Resolving the paradox in high-dimensional space

The identification of ultra-rare cell populations in single-cell transcriptomics is frequently described as a “needle in a haystack” problem. However, our analysis suggests it is more computationally akin to distinguishing a needle made of hay, where rare signals are obscured by the overwhelming covariance of housekeeping genes (Choudhary and Satija, 2022). We have introduced PalmaClust, a framework that redefines the statistical prior for rarity by transposing the Palma ratio— a metric from development economics designed to measure tail inequality—into the domain of gene feature selection.

Unlike the Gini coefficient, which is mathematically most sensitive to changes in the middle of a distribution, the Palma ratio explicitly acts as a high-pass filter for the heavy-tailed distributions characteristic of rare marker genes. By integrating this tail-sensitive metric within a multi-view graph fusion architecture, PalmaClust resolves the fundamental tension in unsupervised learning: the trade-off between preserving global manifold structure and resolving local, high-frequency variation.

### 4.2 Why Gini index fails rare cell types

A critical finding of this study is the specific failure mode of Gini-based feature selection in the presence of background noise. In economics, the Gini coefficient is known to be relatively insensitive to the tails of the income distribution, responding primarily to transfers within the middle class. In transcriptomics, this “middle class” corresponds to housekeeping and metabolic genes with moderate, ubiquitous expression (Eisenberg and Levanon, 2013). Our studies (see **Fig. 2**) demonstrate that as a cell type becomes rarer (*<* 1%), the Gini index fails to prioritize its markers because the signal contribution of the rare tail is mathematically outweighed by technical noise in the abundant middle (Arceneaux *et al*., 2023). In contrast, the Palma ratio (*p*_*top*_*/p*_*bottom*_) effectively discards this stable middle, rendering the metric hyper-sensitive to the “spike-and-slab” expression patterns of rare markers. This confirms that “rarity” in scRNA-seq is not merely a function of low cell count, but of specific distributional inequalities that require specialized statistical treatment.

### 4.3 Multi-view graph fusion

While the Palma ratio provides superior feature selection for rare signals, our results highlight that no single metric can capture the totality of biological heterogeneity. Graphs constructed solely from Palma-selected features tend to fragment major cell lineages, while those based on Fano factor or Gini index preserve global structure but obscure rare cliques. PalmaClust addresses this by fusing these orthogonal topological views. The Palma view injects strong edges between rare cells, while the Fano/Gini views work to maintain global coherence. Empirical benchmarking on the airway dataset (GSE102580) reveals that this fusion allows PalmaClust to occupy a unique position on the Pareto frontier, simultaneously achieving high global stability (ARI > 0.73) and superior rare-cell sensitivity (Ionocyte F1 > 0.86), a balance that single-view methods like GiniClust3 and Seurat fail to achieve.

### 4.4 Biological implications

The capacity to resolve 0.2% populations has profound clinical implications. In the airway epithelium, the clear isolation of pulmonary ionocytes confirms that *CFTR* expression is a rare-cell phenomenon rather than a diffuse mucosal property (Okuda *et al*., 2021). Misclassifying these cells into broad secretory clusters, as observed with standard baselines, would fundamentally mislead gene therapy strategies for Cystic Fibrosis, which rely on precise promoter targeting (Cooney *et al*., 2018). Similarly, in the immune system (GSE94820), the detection of Mono4 (cytotoxic monocytes) highlights the plasticity of myeloid lineages. The Mono4 subset, characterized by NK-like killing machinery, represents a critical effector population in autoimmunity that is frequently averaged out in standard monocyte annotations (Hu *et al*., 2020). By resolving these subsets, PalmaClust ensures that cell atlases reflect the true functional compartmentalization of tissue, rather than a smoothed average.

### 4.5 Limitations and computational constraints

Despite its sensitivity, PalmaClust is subject to specific limitations. The method is highly sensitive to doublets: a hybrid transcriptome formed by two abundant cell types can structurally mimic a rare “intermediate” state (Wolock *et al*., 2019). Rigorous doublet filtration is therefore a mandatory prerequisite. Also, the Palma hyperparameters (*p*_*t*_, *p*_*b*_) may require tuning for detecting populations with varying degrees of rarity, although our default settings proved robust across the tested benchmarks. A future work of consensus clustering based on different Palma settings will be a feasible direction.

